# Exploring respiratory viral pathogens and bacteriome from symptomatic SARS-CoV-2-negative and positive individuals

**DOI:** 10.1101/2024.05.13.593815

**Authors:** Vijay Nema, Sushama Jadhav, Rushabh B Waghmode, Varsha A. Potdar, Manohar Lal Choudhary

**Author notes:** Corresponding author: Dr. Vijay Nema Scientist ‘E’ & Head, Div. of Molecular Biology, ICMR-National Institute of Translational Virology & AIDS Research (formerly ICMR-National AIDS Research Institute), 73 G MIDC, Bhosari, Pune, India 411026.

## Abstract

In the COVID pandemic era, increased mortality was seen despite some unknown etiologies other than SARS-CoV2 viral infection. Vaccination targeted to SARS-CoV2 was successful due to infection caused by pathogens of viral origin based on symptomatology. Hence, it is essential to detect other viral and bacterial infections throughout the initial wave of the COVID-19 disease outbreak, particularly in those suffering from a symptomatic respiratory infection with SARS-CoV-2-negative status. This study was planned to explore the presence of bacterial and other respiratory viruses in symptomatic patients with SARS-CoV2-positive or negative status. The study selected128 patient’s samples out of 200 patients’ samples (100 at each time point) collected for routine SARS-CoV-2 detection schedule in December 2020 and June 2021. Considering the seasonal changes responsible for the occurrence of respiratory pathogens, we finalized 64 SARS-CoV-2 tested patients with 32 SARS-CoV-2-negatives and 32 SARS-CoV-2-positives from each collection time to examine them further using real-time PCR for the presence of other viral species and bacterial infection analyzing 16S rRNA metagenome supporting to cause respiratory infections. Along with various symptoms, we observed the co-infection of adenovirus and influenza B(Victoria) virus to two SARS-CoV-2-positive samples. The SARS-CoV-2-negative but symptomatic patient showed Rhinovirus (7/64 i.e. 10.9%) and Influenza (A/H3N2) infection in 4 patients out of 64 patients (6.25%). Additionally, one SARS-CoV-2-negative patient enrolled in June 2021 showed PIV-3 infection. Influenza A/H3N2 and Adenovirus were the cause of symptoms in SARS-CoV-2-negative samples significantly. Thus, the overall viral infections are considerably higher among SARS-CoV-2-negative patients (37.5% Vs 6.25%) compared to SARS-CoV-2-positive patients representing respiratory illness probably due to the abundance of the viral entity as well as competition benefit of SARS-CoV-2 in altering the imperviousness of the host. Simultaneously, 16S rRNA ribosomal RNA metagenomenext-generation sequencing (NGS) data from the same set of samples indicated a higher frequency of Firmicutes, Proteobacteria, Bacteroidota, Actinobacteriota, fusobacteriota, Patescibacteria, and Campilobacterotaphyla out of 15 phyla, 240 species from positive and 16 phyla, 274 species from negative samples. Exploring co-infecting respiratory viruses and bacterial populations becomes significant in understanding the mechanisms associated with multiple infecting pathogens from symptomatic COVID-positive and negative individuals for initiating proper antimicrobial therapy.

**Author Summary:** Frequent transfer of SARS-CoV-2 events has resulted in the emergence of other viral infections along with several evolutionarily separate viral lineages in the global SARS-CoV-2 population, presenting significant viral variants in various regions worldwide. This variation also raises the possibility of reassortment and the creation of novel variants of SARS-CoV-2, as demonstrated by the COVID pandemic in all the waves, which may still be able to cause illness and spread among people. Still unclear, though, are the molecular processes that led to the adaption of other viral and bacterial pathogens in humans when a human SARS-CoV-2 virus was introduced. In this study, we identified the presence of various other viral infections and bacterial content in symptomatic COVID-19-positive and negative patients, as evidenced by the data obtained using next-generation sequencing of 16S rRNA metagenome and real-time PCR detection technologies. Symptoms might have been induced by bacterial content and various viral entities other than the SARS-CoV-2 viral infection in the COVID-negative population, indicating its importance in detecting and initiating appropriate therapy to recover from all other infections.

## Introduction

After its first detection in Wuhan, China, the SARS-CoV-2 was found to spread rapidly to other parts of China and beyond and become a global threat[1]. By April 10, 2023, the pandemic pathogen, Severe Acute Respiratory Syndrome Corona virus 2, abbreviated as SARS-CoV-2, had infected 684 million people and caused 6.8 million deaths worldwide through 657 million cases recovered from the infection [2]. Nations, including the USA, India, France, Germany, Brazil, Japan, South Korea, Italy, the UK, and Russia, were the most affected globally. The United States reported the highest number of identified cases in a single country at over 105 million and over 1148 thousand deaths, followed by India with more than 44 million cases and over 530 thousand deaths.

A focus on respiratory viral pathogens and bacteriome gained attention in various studies throughout the COVID-19 pandemic, and the number of publications addressing the entire microbial community in this area has grown significantly [3]. The pandemic was observed due to the activation of predominant species of various respiratory viral pathogens and bacteriome in nasopharyngeal samples. However, there was the presence of multiple such microorganisms in healthy individuals, as reported earlier by De Boeck et al., wherein they proposed that the nasopharyngeal bacteriome of individuals in good health was divided into four profiles with the first type of bacteria being Moraxella, followed by Fusobacterium, Streptococcus, and a combination of Corynebacterium, Staphylococcus, and Dolosigranulum. However, with NGS analysis, we found numerous other bacterial populations in SARS-CoV-2 positive and symptomatic negative individuals in the COVID-19 pandemic era [4].

In India, the Maharashtra state experienced the highest positive cases, followed by many states in South India [5]. We collected samples for the present study in December 2020 and June 2021, when 731629 and 2280282 COVID patients were reported in India. These were among the most reported months, with two different weather conditions. Surprisingly, COVID-19 is just one of the pathogens among the respiratory viruses listed; so far, it has made us understand the threat of respiratory viruses and the impact of respiratory tract infections. Research Interest has increased over the past decade due to its critical involvement in the growth of numerous ailments, including recurrent respiratory conditions [6].

Next-generation sequencing (NGS) has recently been considered a powerful diagnostic platform for detecting pathogens in symptomatic COVID-19-positive and negative patients [7]. The conception of unprejudiced sequence analysis of 16S rRNA metagenome from nasopharyngeal swabs allowed us to identify prevalent bacterial population in one single test, including Gram-positive microorganisms like Firmicutes and Actinobacteria species that cannot be cultured without any antimicrobial treatment. Very few publications have shown co-infections amongst SARS-CoV-2 and other airborne pathogens capable of producing common clinical manifestations, even though those other airborne pathogens became the focus of extensive investigation in the period before SARS-CoV-2 arose [8,9]. Tracking infecting pathogens is just one of many approaches that research facilities need to prioritize and modify during this epidemic era due to the observance of enormous clinical findings [10,11]. Identifying existing viral pathogens in circulation might assist in the development of scientifically proven tactics to combat the outbreak in the upcoming flu seasons and subsequent epidemic spikes as an enormously growing percentage of individuals require testing for COVID-19-like symptoms. A wide variety of clinical symptoms of respiratory infections are represented along with the symptoms of COVID-19, illustrating moderate respiratory disease to severe pneumonia. Practical lab tests are crucial to differentiate between genuine COVID-19 cases and starting public health measures to stop the further transmission of SARS-CoV-2 [12]. Considering the seasonal variation in the symptoms during the second wave of COVID-19 in India, many more variants of this emerging viral strain like B.1.1.7 (Alpha), B.1.351 (Beta), and B.1.1.28.1 (Gamma), B.1.1.28.2 (Zeta), B.1.617.1 (Kappa) and B.1.617.2 (Delta) with its derivatives were also circulating in the majority of patients from various regions [13]

Critical monitoring of various co-infecting viral pathogens and bacterial populations representing the same symptoms as shown during the SARS-CoV-2 virus infection, ultimately leading to COVID-19, is vital for developing proper treatment modalities. As reported earlier, the Indian population is vulnerable to multiple infections with variable frequencies, seasonal changes, and geographic locations. Hence, the current study aimed to explore the presence of other viral and bacterial entities in symptomatic patients who were positive and harmful for SARS-CoV-2 infection. We utilized the previously optimized protocols for identifying other viruses using RT-PCR and bacterial populations using next-generation sequencing technology (NGS), focusing on 16S rRNA metagenome analysis.

## Results

Nasopharyngeal samples were chosen to cover the most sensitive age for infection as reported through this pandemic, with age groups ranging from 18-35 with a mean age of 27.5. The number of male patients was 41(64%) and female patients were 23(36%). The majority of the patients detected other respiratory viruses from COVID-negative status, having Rhinovirus (HRV) (10.9%) 7/64 followed by INF (A/H3N2) (6.25%)4/64. Co-infection with adenovirus and influenza B (Victoria) was found in two SARS-CoV2 positive samples, one each from two collection periods. The most expected observations presented by patients were fever, cough, nasal discharge, and chest congestion, which are described in Table 1. Out of 64, fourteen (21.87%) patients harbored other viral infections.

**Table 1:** Demographic details of the patients along with symptoms and types of other respiratory viruses detected.

| Patient ID<br>Dec 2020 | Gender | Age | Symptoms | Virus detected |
| --- | --- | --- | --- | --- |
| P13 | F | 21 | Fever, Body ache | SARS-CoV-2, Adenovirus |
| P38 | M | 26 | Body ache | SARS-CoV-2, INF-B(VICTORIA) |
| N7 | F | 34 | Cough, Nasal discharge | Rhinovirus |
| N9 | F | 30 | Sore throat | Rhinovirus |
| N11 | M | 33 | Cough, Nasal discharge | Rhinovirus |
| N14 | M | 30 | Sore Throat, nasal discharge | Rhinovirus |
| N22 | M | 27 | Sore throat | INF-A/H3N2 |
| N23 | M | 23 | nasal discharge | Rhinovirus |
| N25 | M | 30 | Fever, Cough, Nasal discharge | INF-A/H3N2 |
| N32 | F | 28 | Body ache | Rhinovirus |
| N48 | M | 28 | cough, fever | INF-A/H3N2 |
| N54 | M | 23 | cough, fever, Sore Throat | Rhinovirus |
| N40 | F | 18 | Fever | PIV3 |
| N47 | F | 21 | cough, fever | INF-A/H3N2 |

### Analysis of samples for other viral infections in the winter season

In the samples collected in December 2020, most of the positive patients showed signs and symptoms like nasal discharge (55%), fever of more than 100°F (high-grade fever) (22.2%), cough (33%), and sore throat (33%). The majority of HRV-positive patients had nasal discharge as the primary symptom. In a group of SARS-CoV-2 positive patients (n=32), 13 patients (40.62%) were female, while 19 (59.37%) were males. In these SARS-CoV-2 positive patients, the highest numbers of patients were having body aches (25%) with other symptoms such as cough (18.75%), nasal discharge (9.3%), fever (9.3%), sore throat (9.3%), Breathlessness (3.12%), Nausea (3.12%), etc. In SARS-CoV-2 negative patients (n=32), there were 22 (68.75%) males, while 10 (31.25%) were female patients. The dominant symptom in these patients was sore throat (43.75%), which might have raised the alarm in the panic era. While other noted symptoms in these patients were fever (34.37%), nasal discharge (25%), cough (21.87%), Body ache (15.6%), Breathlessness (6.25%), Nausea (3.12%), etc. The observation of clinical symptoms from SARS-CoV-2 positive and negative patients collected in Dec 2020 indicated higher symptoms are presented in Figures 1 A and B, respectively.

**Figure 1:**
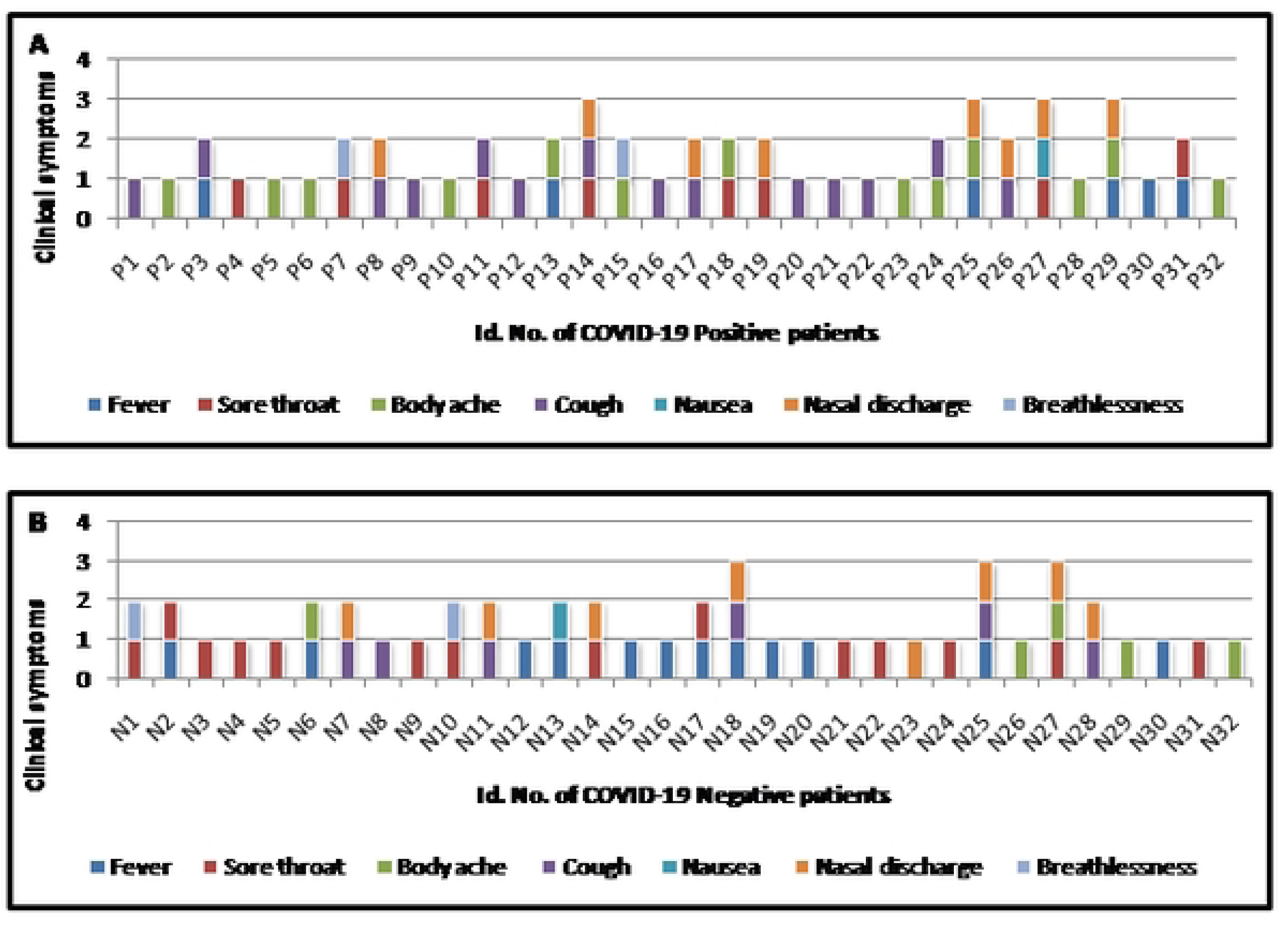
Clinical symptoms of A) COVID-19 positive patients B) COVID-19 Negative patients during Dec 2020 (winter season). Each colored vertical box in the figure represents about a symptom noted during enrollment of the patient for suspected SARS-CoV-2 infection.

### Analysis of samples for other viral infections in the summer season

In the samples collected in June 2021, most of the positive patients showed signs and symptoms like a fever of more than 100°F (high-grade fever) (80%), cough (60%), body aches (20%), and sore throat (20%) while no patient found with nasal discharge in this period compared to samples collected in December 2020. The observation of clinical symptoms from SARS-CoV-2 positive and negative patients collected in June 2021 indicated fewer symptoms, as presented in Figure 2 A and B, respectively, due to seasonal changes (temperature and humidity). Additionally, the clinical symptoms of patients with co-infection and other viral infections are presented in Figure 3. Furthermore, the percentages of various viruses detected from study samples are presented in Figure 4, which depicts the prevalence of multiple viruses reported during the collection period of the study samples.

**Figure 2:**
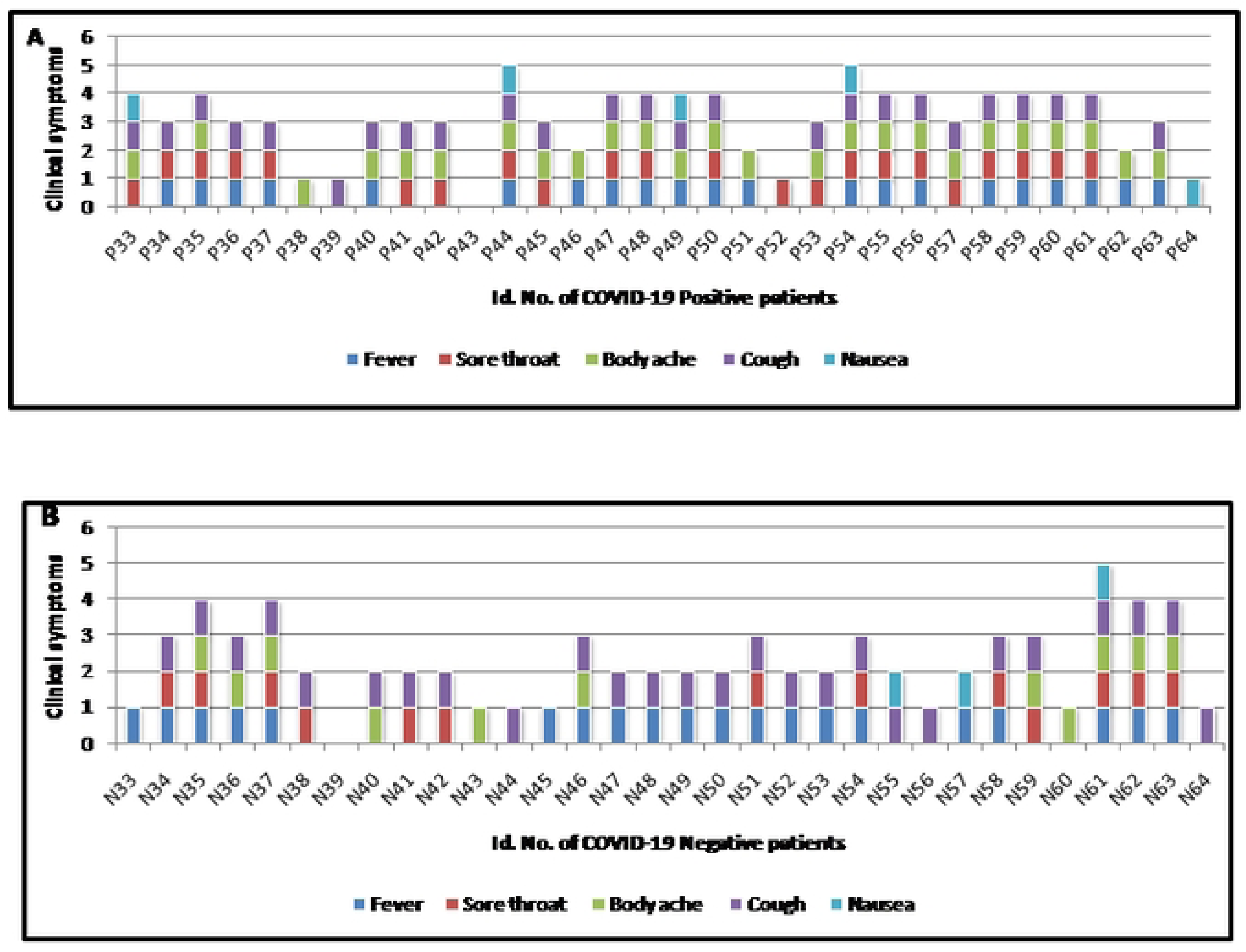
Clinical symptoms of A) COVID-19 positive patients B) COVID-19 Negative patients during June 2021 (summer season). Each colored vertical box in the figure represents about a symptom noted during enrollment of the patient for suspected SARS-CoV-2 infection.

**Figure 3:**
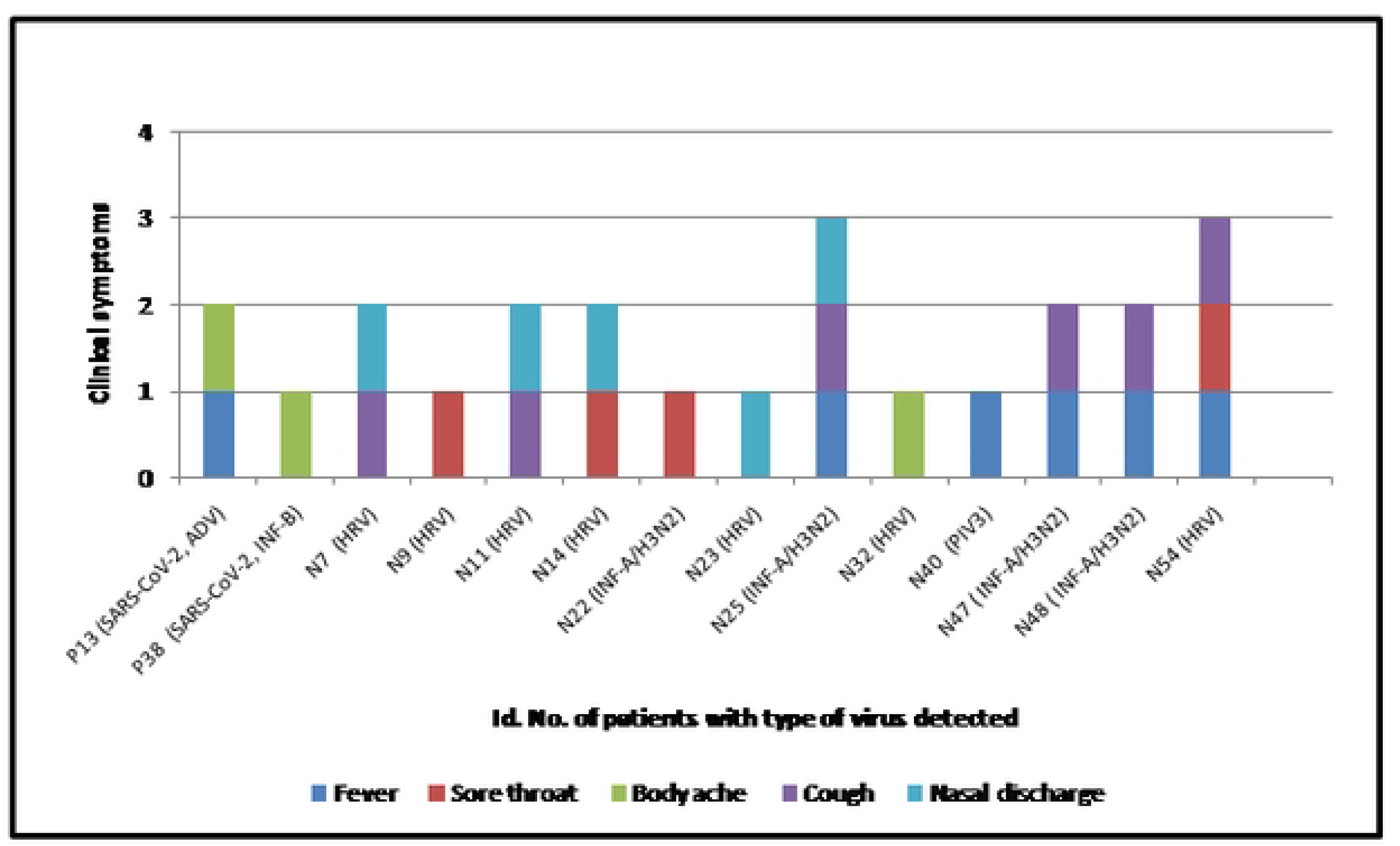
Clinical symptoms of patients with co-infection and other viral infection. Each colored vertical box in the figure represents about a symptom noted during enrollment of the patient for suspected SARS-CoV-2 infection.

**Figure 4:**
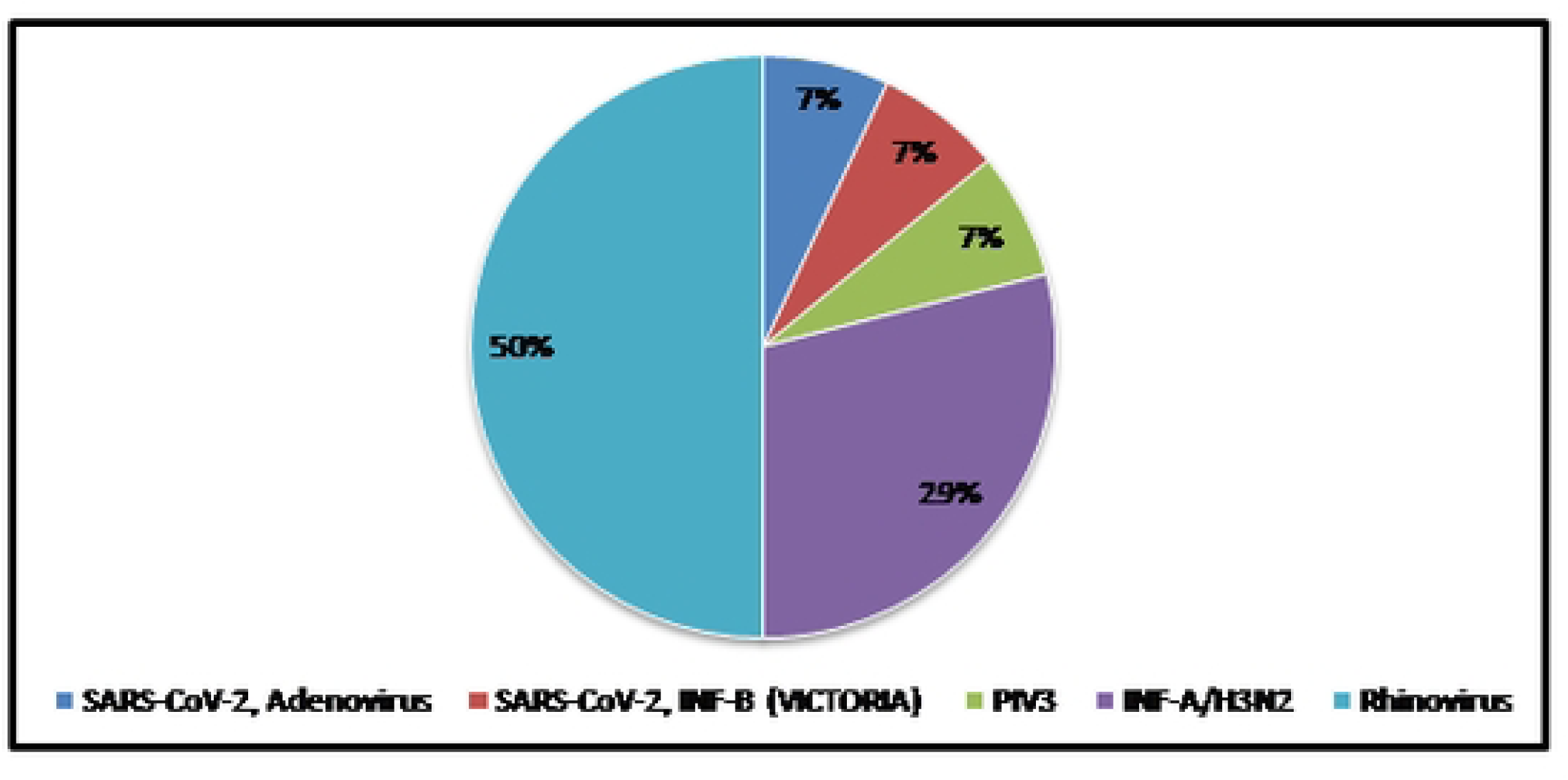
Percentages of co-infecting and other viral infections from study samples

### Analysis of 16S rRNA metagenome sequences

Bacterial communities were analyzed by applying 16S rRNA (ribosomal RNA) genetic material amplification to the next-generation sequencing process. The results showed that taxonomic diversity was less in COVID-19-positive patients compared to COVID-negative patients, and microbial alpha-diversity indices were not significantly different among the two groups of patients. These nasopharyngeal swabs showed the maximum frequency of phyla like Fusobacteriota, Actinobacteriota, Bacteroidota, Firmicutes, Proteobacteria, Campilobacterota, and Patescibacteria in COVID-negative samples. In contrast, Fusobacteriota, Actinobacteriota, Campilobacterota, Bacteroidota, Firmicutes, Proteobacteria, and Patescibacteria in COVID-positive samples that accounted for almost 50% of all the sequences characterizing the communities of microbes in both the groups Figure 6A, B, C. The Relative frequency of occurrence of bacteria at taxonomic level was determined for COVID-19-positive (n=32) and -negative (n=32) nasopharyngeal samples. An average frequency of sequences per taxa was calculated from the total sum of all sequence counts, and the standard deviation (SD) is shown in percentage (%). The taxonomic data is included for phylum where its occurrence is > 1%, as depicted in Table 3. Furthermore, the compositions to determine the diversity of 16S rRNA metagenome for COVID-negative and positive samples are explored using the Krona chart, as shown in Figure 7A and 7B, respectively. The krona chart is a multilayer pie chart for Positive and Negative samples representing complex high-level categorization into specialized sections towards the exterior of the round, displaying rankings in a more dynamic manner with its percentage predominance. This chart helps to display larger phyla in greater depth and clarity while grouping and labeling smaller phyla

**Figure 6A:**
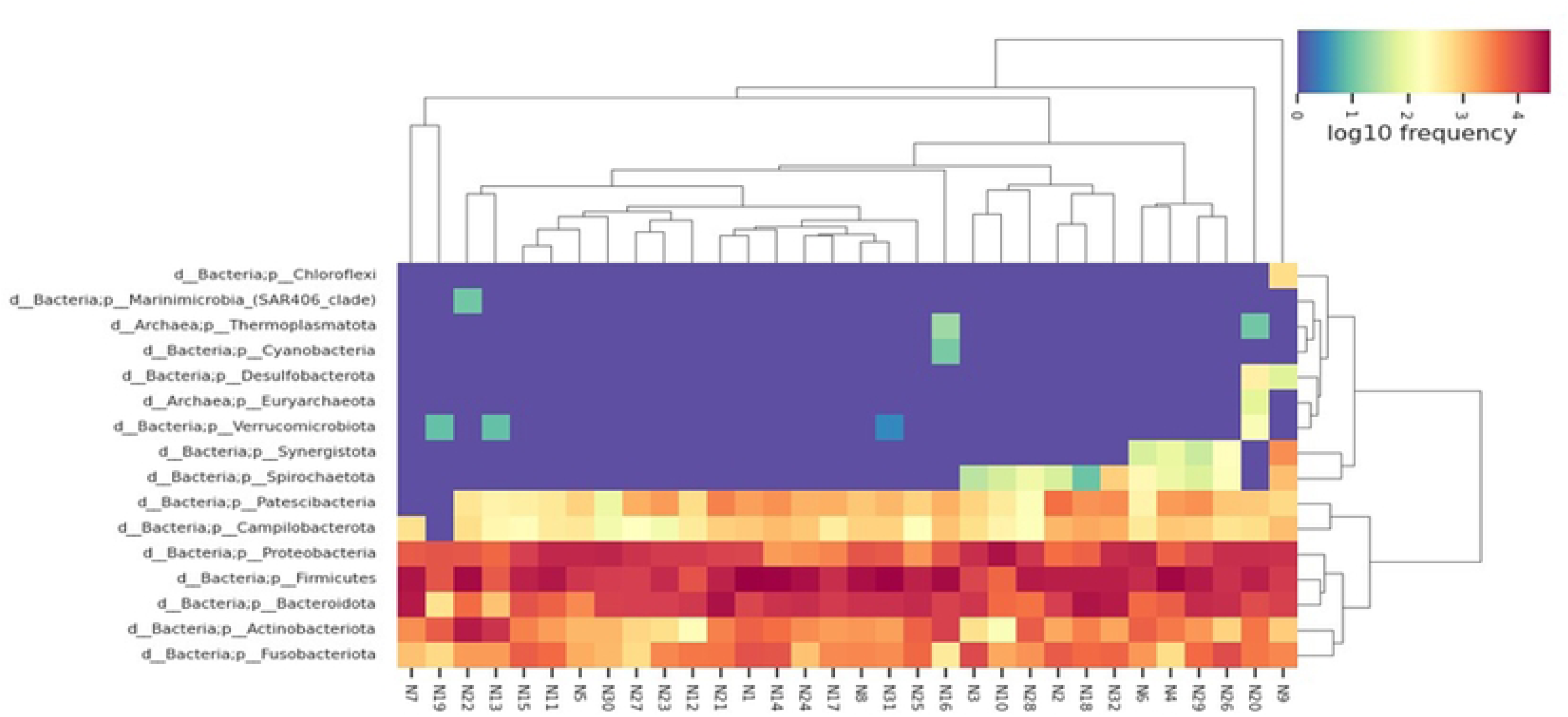
Heatmap showing the 16 phyla, 168 Genera & 274 species found in the negative samples. These nasopharyngeal swabs showed the maximum frequency of phyla like Fusobacteriota, Actinobacteriota, Bacteroidota, Firmicutes, Proteobacteria, Campilobacterota, and Patescibacteria. There was the presence of bacterial variety at each site. The red bar indicates a relatively high abundance of approximately 90% while orange-yellow blocks and green, purple blocks represent relatively medium and low frequencies respectively of these phyla.

**Figure 6B:**
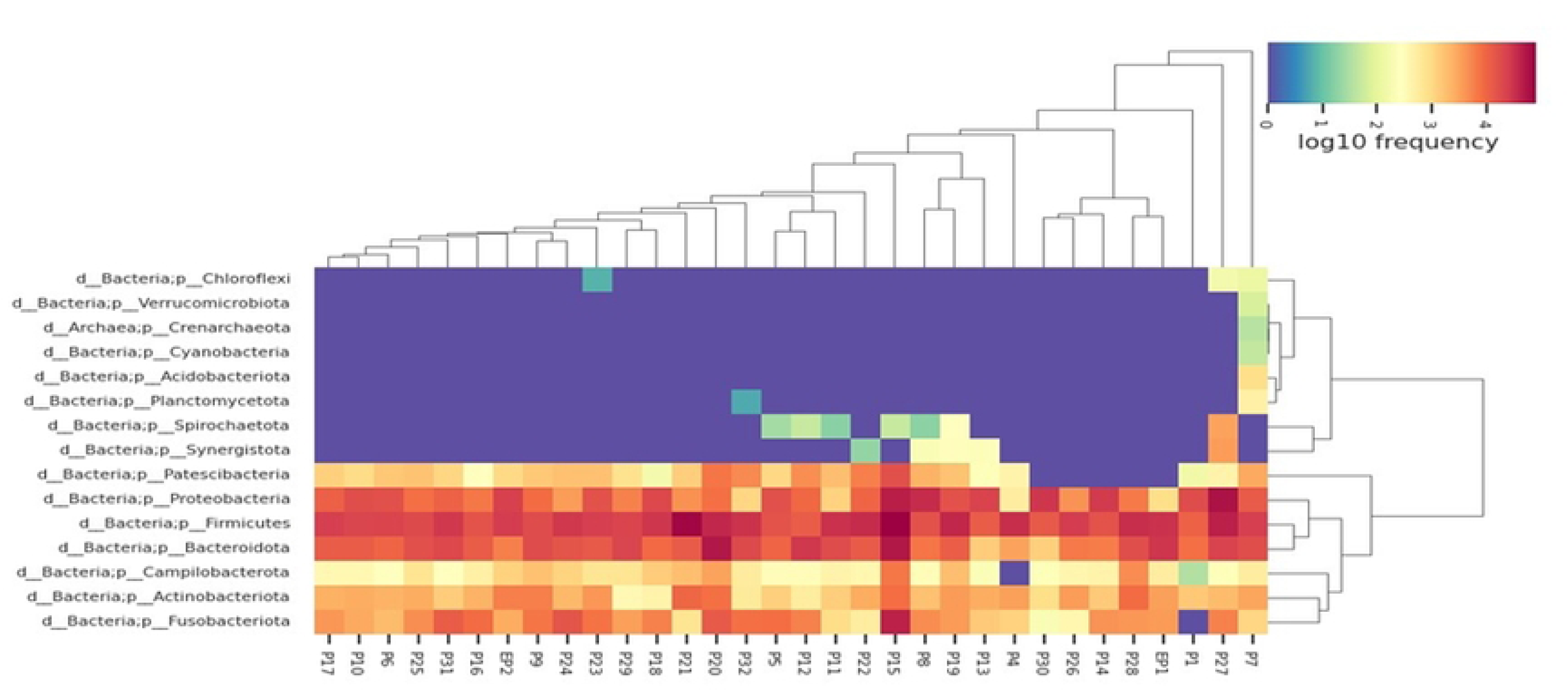
Heatmap showing the 15 phyla, 145 Genera & 240 species found in the positive samples. These nasopharyngeal swab samples showed maximum frequency of phyla like Fusobacteriota, Actinobacteriota, Campilobacterota, Bacteroidota, Firmicutes, Proteobacteria, and Patescibacteria along with bacterial variety at each site. The red bar indicates a relatively high abundance of approximately 90% while orange yellow blocks and green, and purple blocks represent relatively medium and low frequencies respectively of these phyla.

**Figure 6C:**
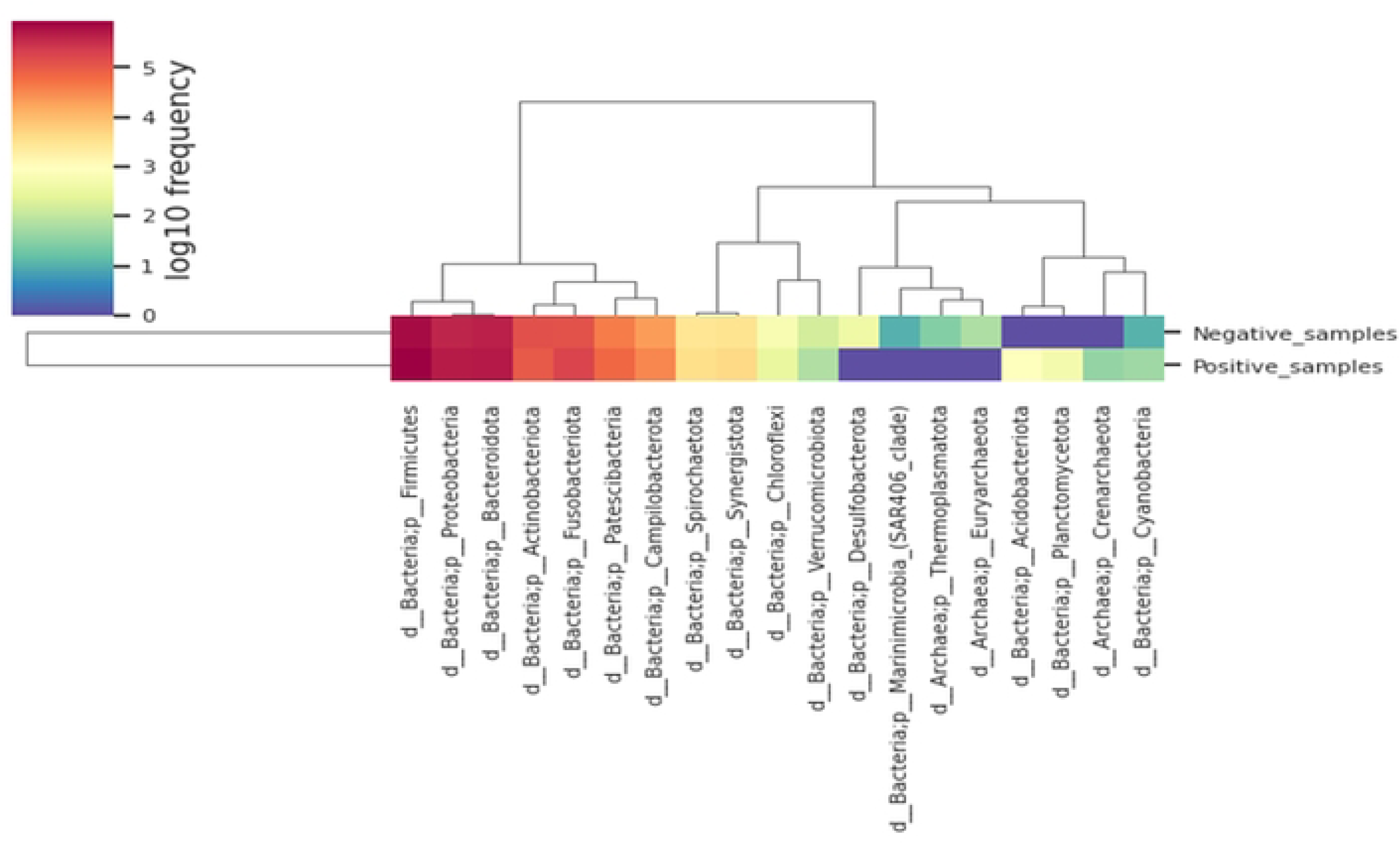
Heatmap showing the 19 phyla commonly observed in both positive and negative samples. These nasopharyngeal samples had shown maximum frequency of phyla like Firmicutes, Proteobacteria, Bacteroidota, Actinobacteriota, fusobacteriota, Patescibacteria and Campilobacterota with bacterial variety at each site. The red bar indicates relatively high abundance of approximately 90% while yellow blocks and green, purple blocks represent relatively medium and low frequency respectively of these phyla.

**Figure 7A:**
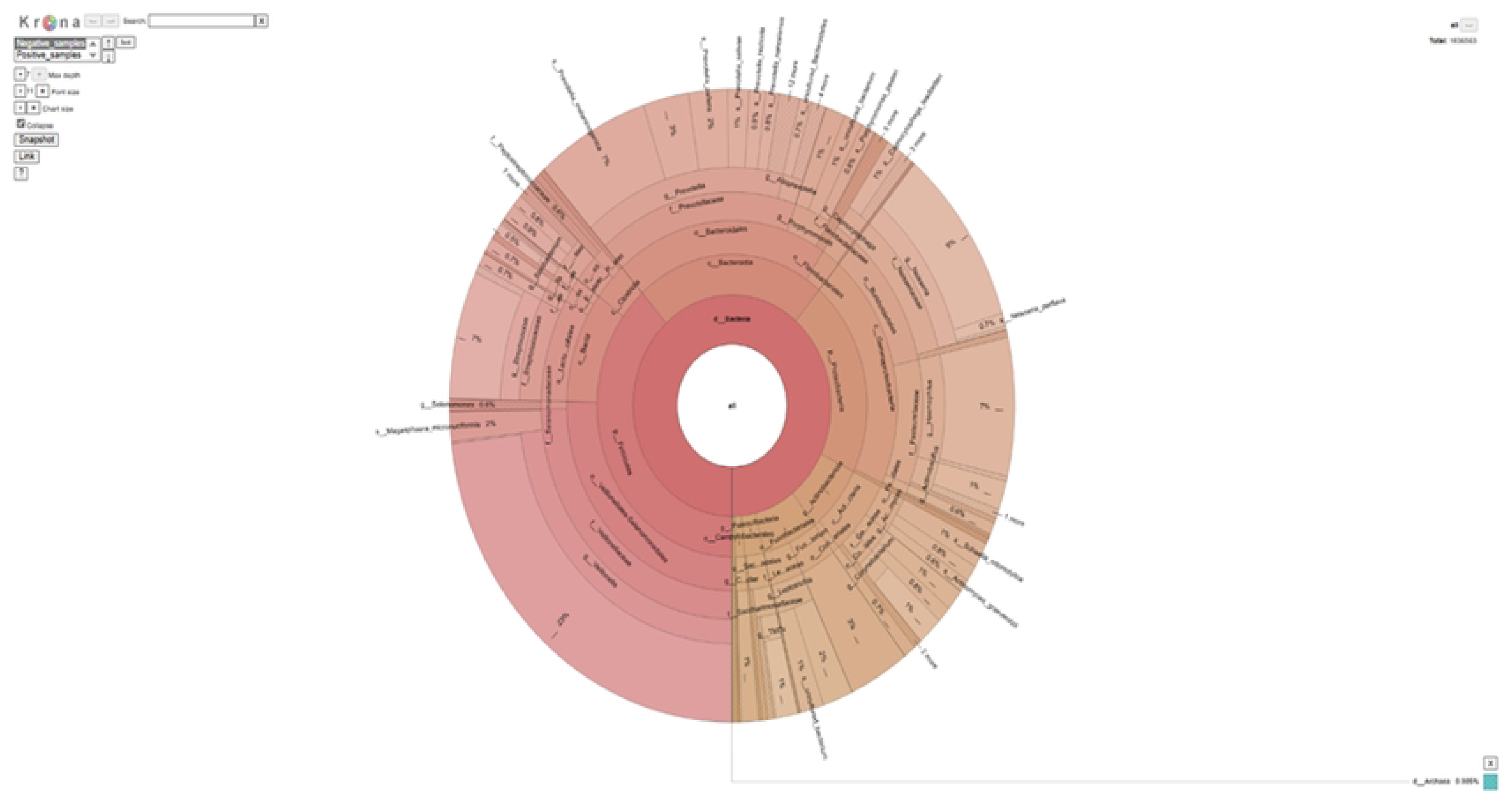
The multilayer pie chart for Negative samples representing complex high-level categorization into specialized sections towards the exterior of the round, displaying rankings in a more dynamic manner. This chart helps to display larger phyla in greater depth and clarity while grouping and labeling smaller phyla.

**Figure 7B:**
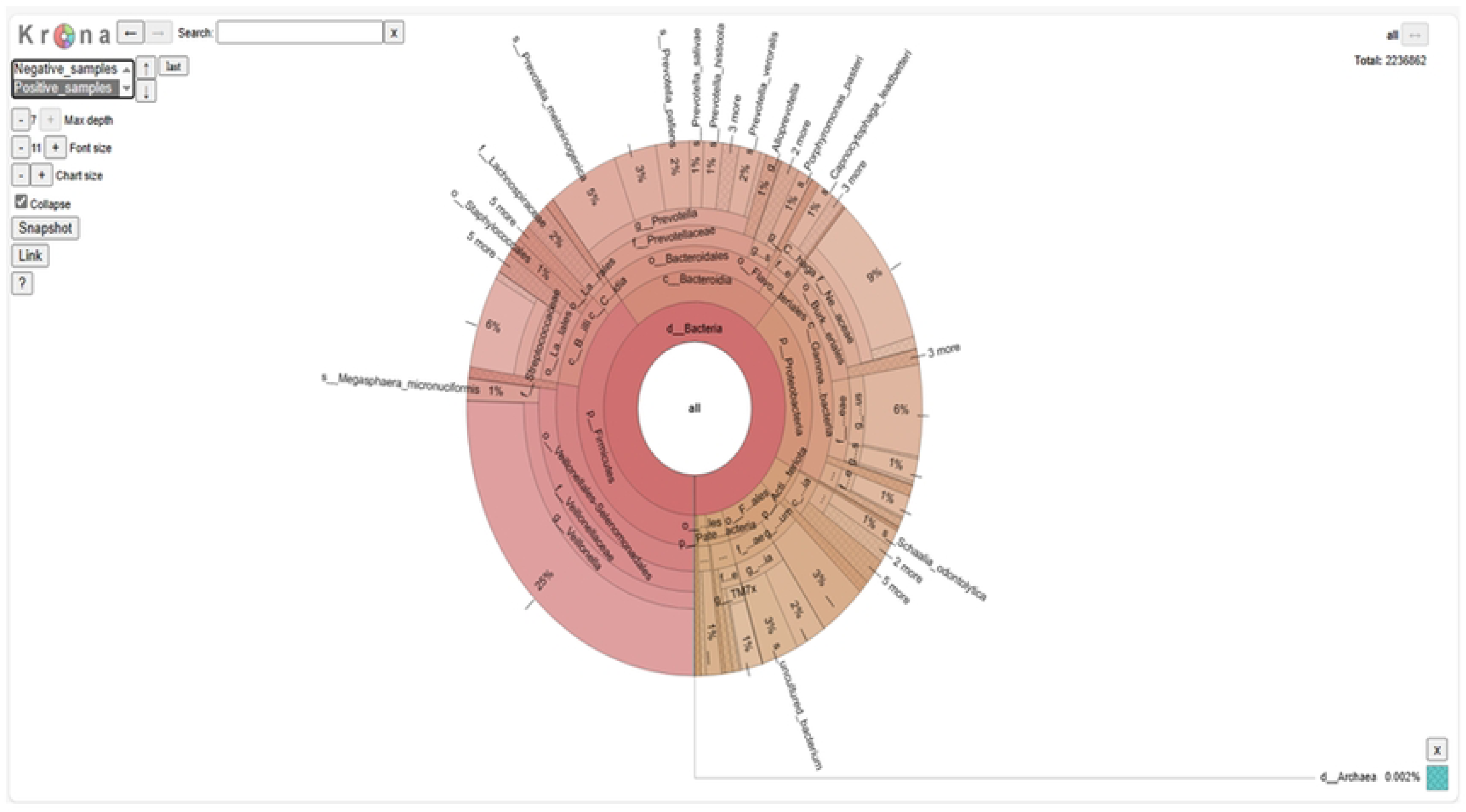
The multilayer pie chart for Positive samples representing complex high-level categorization into specialized sections towards the exterior of the round, displaying rankings in a more dynamic manner. This chart helps to display larger phyla in greater depth and clarity while grouping and labeling smaller phyla.

## Discussion

The COVID-19 pandemic started with the suspicion of SARS-CoV-2 infection in many patients suffering from acute upper or lower respiratory tract infections. Influenza virus (IAV, H1N1, IBV), parainfluenza virus (HPIV), human metapneumovirus (HMPV), human Bocavirus (HBoV), Human respiratory syncytial virus (HRSV), adenovirus (HAdV), Rhinovirus (HRV), enterovirus (EV), and corona viruses constitute the most often associated viruses in acute respiratory infections (ARI) [14,15]. Older people having related illnesses, including coronary artery disease, hypertension, diabetes mellitus, chronic kidney disease, immunosuppressive people, etc., have been linked to the severity of the infection and an elevated death rate. Although molecular techniques have been developed practically everywhere in various countries, single-plex PCR is time-consuming and challenging for detecting numerous pathogens in one attempt. Hence, using a multiplexing approach in the Real-time PCR technique, we determined the frequency of other viruses present among study patients during the COVID-19 pandemic. Early detection is crucial for prompt treatments and the start of a precise antimicrobial/antiviral treatment. A correct diagnosis would help these people to take unwanted medicines, which would slow the development of pathogens that spread respiratory infections. Medication of people who have a quick assessment and definitive therapeutic interventions with the multiplexing diagnostic facility for COVID-19-positive patients is essential throughout this time for improved treatment of patients.

Interestingly, it is observed from this pilot study that a more significant number of the SARS-CoV-2 negative male patients indicated the presence of other viral infection (Rhinovirus, IF-A(AH3)) compared to female patients from the same group and all SARS-CoV-2 positive patients. Additionally, a comparative observation of the symptoms of patients collected at two different periods indicated an absence of nasal discharge in patients whose samples were collected in June 2021 compared to samples collected from patients in December 2020. The SARS-CoV-2-positive patients showed signs and symptoms like nasal discharge (55%), high-grade fever (22.2%), cough (33%), and sore throat (33%). The majority of HRV-positive patients had nasal discharge as the primary symptom. In a group of SARS-CoV-2 positive patients (n=32), 13 patients (40.62%) were female, while 19 (59.37%) were males. In these SARS-CoV-2 positive patients, the highest numbers of patients were having body aches (25%) with cough (18.75%), nasal discharge (9.3%), fever (9.3%), sore throat (9.3%), Breathlessness (3.12%), and Nausea (3.12%). In SARS-CoV-2 negative patients (n=32), there were 22 (68.75%) males, while 10 (31.25%) were female patients. Observance of sore throat was the predominant symptom (43.75%), alarming the situation in the COVID-19 pandemic era. Additionally, patients were having fever (34.37%), nasal discharge (25%), cough (21.87%), Body ache (15.6%), Breathlessness (6.25%), and Nausea (3.12%).

Cong et al., 2022 reported that antibiotic use and bacterial infection for hospitalized ARI patients during the COVID-19 pandemic were variable concerning variable time points. Staphylococcus aureus has been the commonly documented conferring resistance organism in COVID-19 patients, followed by Pseudomonas aeruginosa, Escherichia coli, and Klebsiella pneumonia. The cumulative documented microbial risk of transmission in COVID-19 patient populations was 10.5%[16]. According to the current scenario, most COVID-19-positive patients did not show much co-infection with other viral pathogens, which is consistent with research articles published earlier[17]. The competitiveness of SARS-COV-2 in regulating the human immune system is one potential explanation for such a scenario[18].

The widespread use of Illumina sequencers greatly aids many sequencing projects. NGS technologies are widely used for a variety of purposes. Initially, these technologies provide remarkable precision in sequencing, guaranteeing dependable outcomes. Furthermore, the price per gigabyte (Gb) of data acquired using these techniques is low. Additionally, various equipment alternatives are available on the market, enabling researchers to choose the best instruments to suit their project demands. Sequencing machinery ranges from small, moderately performing benchtop devices like MiniSeq to huge, population-based initiative-specific devices like HiSeqX used to sequence complete genomes. Healthcare professionals can take immediate action based on the type of disease to avoid or limit its large-scale spread due to the ability of next-generation sequencing to rapidly identify pathogens that pose a hazard to public health. NGS is, therefore, essential for the clinical diagnosis of infections. In most places, researchers used whole genome sequencing and single nucleotide polymorphisms in clinical outbreaks. Isolates of methicillin-resistant S. Aureus are compared, giving medical professionals quick access to clinically relevant validated data[19]. Additionally, NGS is a potent technique for determining a pathogen’s virulence factors, which might enhance public health interventions.

Furthermore, NGS can help prevent infections by aiding the avoidance of large-scale pathogen outbreaks through the phylogenetic analysis of isolates. Using a neighbour-joining tree or maximum likelihood tree for phylogenetic research can assist in identifying the branch on which the sequences have been clustered [20]. Hence, NGS technologies allow for high-throughput sequencing of DNA, providing massive amounts of sequence data where DNA is fragmented, sequenced in parallel, and then bioinformatically analyzed to identify microbial DNA. Thus, NGS is powerful for metagenomics analysis, allowing the detection of diverse microbial species in complex samples.

The current study outcome indicated that the most prevalent pathogens, along with or without SARS-CoV-2 infection, are HRV and IAV. Due to the confirmed identification of other viral infections, it might be possible to avoid using antibiotics, which would prevent the emergence of resistant bacteria that spread various illnesses. In the future, multiplexing assays will be essential to provide better medical care to patients with earlier diagnoses of other viral and co-viral infections for conclusive treatment. Altogether, we could identify a few drawbacks of this study, which indicated the presence of very few other viral infections and co-infections due to the small number of samples collected from a single hospital.

## Material and methods

### Ethics statement

The samples selected from routine COVID testing to carry out this study were approved by the Institutional Ethics Committee, with permission for the patient to waive consent, as the data was collected anonymously by a non-study institutional committee. The approval number is NARI/EC/Approval/20-21/437, dated December 15, 2020.

### Nasopharyngeal swab sample collection

Earlier studies have demonstrated the viability of utilizing nasopharyngeal swab VTM collection for microbiota investigation and SARS-CoV-2 identification where the nasopharyngeal swabs (NPS) were collected in compliance with CDC regulations. Following these regulations, NPS for diagnostic testing were collected using clean synthetic-head, plastic-shaft swabs, and they were then put into appropriately labelled collecting containers holding three milliliters of VTM.

### Nasopharyngeal swab (NPS) sample processing

The NPS samples were sent to the National AIDS Research Institute from the local Municipal Corporation Hospital, Bhosari, Pimpri-Chinchwad, Pune, for routine testing of SARS-CoV-2. The study selected a total of 128 patient samples out of 200 samples (100 at each time point) collected for routine SARS-CoV-2 detection in December 2020 (winter) and June 2021(summer). The purpose of selecting two different periods was due to weather conditions influencing the infection rate of other viral entities and SARS-CoV-2 infection. We have chosen 64 patient samples from December 2020 and June 2021 from this pool. Two swabs were collected, each from nasal and pharyngeal sources, in a single tube using a Viral Transport Medium (VTM). The data provided along with the samples was anonymized for information about the patient’s age, gender, and symptoms and the SARS-CoV-2 results were made available by the institutional data management unit for SARS-CoV-2 diagnosis at ICMR-NARI.

In the current study, we selected 128 nasopharyngeal samples collected in two seasons and tested for SARS-CoV-2 infection. Out of 128 samples, 64 samples were selected from two different time points with the symptomatic status of the patient consisting of 32 positives for SARS-CoV-2 and 32 SARS-CoV-2 negative as per our routine diagnostic results of real-time PCR. We finalized 32 samples each from SARS-CoV-2 negative (labelled N1-N32; n=32/64) and positive samples (labelled P1, P4-P32, EP1, EP2; n=32/64) having symptoms of LRTI/URTI as suspected for SARS-CoV-2 in two batches and examined the same for the presence of other viral species that causes respiratory infections. Finally, this pilot study considered only 64 samples; the rest were not analyzed. Mainly, the patients with various symptoms included spasms, rhinorrhea, and high fever with the suspected infection of SARS-CoV-2. From both slots, selected samples were in the age group ranging from 18-35 years, and the mean age was 29.34 years in negative patients while 29.84 years in positive patients tested for SARS-CoV-2 infection.

### Isolation of RNA and detection of SARS-CoV-2 using Real-Time PCR

Initially, the SARS-CoV-2 viral RNA was extracted from the NPS samples collected in VTM with the help of ready-to-use kits developed by Meta Design Solutions, Gurgaon, India, followed by testing of SARS-CoV-2 viral infection using COVIDSURE kit developed by Trivitron Healthcare Lab systems Diagnostics, Chennai, India with the help of a multiplex platform of real-time PCR (Model CFX96, Bio-Rad, Hercules, CA, USA). This multiplex assay was performed in a single tube, and E gene expression was considered for the screening test. In contrast, ORF and RdRP gene expression were used to confirm the SARS-CoV-2 infection, and beta-actin gene expression was considered a housekeeping gene. Finally, the ORF expression was taken into account when analyzing Ct values. Considering the threshold cycle (Ct) value for identifying predefined genes for the viral genes examined was below 35, the outcome was deemed positive.

### Detection of other viral strains by multiplex qRT-PCR

RNA isolation was done from all the NPS samples using a Zybio nucleic acid extraction kit following the manufacturer’s protocol (Zybio Inc., China). Using duplex real-time reverse-transcription-polymerase chain reaction (qRT-PCR), around 13 other respiratory viruses and influenza viruses were searched in all the clinical samples using a protocol described earlier [21]. All the clinical samples were tested for the viruses in this study viz: influenza A and subtypes (A/H1N1, pdm09, and A/H3N2), influenza B along with housekeeping RNaseP gene (CDC, WHO), Respiratory syncytial virus (RSV) A & B, Human metapneumovirus (hMPV), Parainfluenza (PIV) virus 1, 2, 3, 4, Rhinovirus and adenovirus, along with human corona virus OC43, 229E, NL63, HKU1 [22]. Briefly, the viral RNA was considered for first-strand cDNA synthesis following the protocol mentioned in the one-step RT PCR kit (SuperScript™ III kit, Invitrogen, USA) consisting of various reagents from the kit inclusive of 10 μmol of each primer, 2X buffer (12.5 μl), 0.5μl SuperScript™ III enzyme, 5μlnucleic acid and 5μmol of TaqMan probe. The programming conditions of real-time PCR were defined in the manufacturer’s protocol book. The data based on fluorescence was collected at different steps of the programming conditions, and the threshold value was interpreted at the end of the assay. [21].

### Isolation of bacterial DNA and 16S metagenome sequencing

Using the protocol published by Qiagen to isolate bacterial DNA using the DNeasy UltraClean Microbial kit, the bacterial DNA was isolated from NPS samples (200 µl) collected in VTM. The bacterial DNA was further processed for a two-stage PCR procedure with a PCR-next-generation sequencing (NGS) strategy to characterize the 16S rRNA V3-V4 metagenome[23]. The V3-V4 variable region of microbial 16S rRNA metagenomes was amplified using specially designed primers for Illumina sequencers. The PCR reagent blanks as negative controls were amplified and sequenced along with the NPS samples. An Illumina MiniSeq with a mid-output kit and paired-end 155 base reads were used for the sequencing process and analyzed subsequently [24].

### Analysis of 16S rRNA metagenomic sequences

The PEAR (Paired-End read merger) technique (v0.9.11) was utilized to merge raw sequences acquired from the sequencer. Combined sequences within the QIIME2 (v 2020.8.0) workflow were quality-filtered and denoised using the DADA2 algorithm [25]. For all subsequent analyses, the amplicon sequence variations (ASVs) were created and used further for downstream evaluation. The naïve Bayes taxonomy classifier was used to assign taxonomy to ASVs using the Bayes taxonomy classifier trained with the SILVA SSU 138 OTU database [26]. Alpha diversity was calculated using a rarefaction curve estimated for 5000 sequences per sample that related the growth of several species revealed as a function of individual samples (Figure No. 5). The data indicating Shannon Index, Simpson’s Index, observed data richness and evenness were measured at ASV level. The mean index score and the standard deviation (SD) were calculated for COVID-19-negative (n = 32) and COVID-19-positive (n = 32) samples as presented in Table 2.

**Figure 5:**
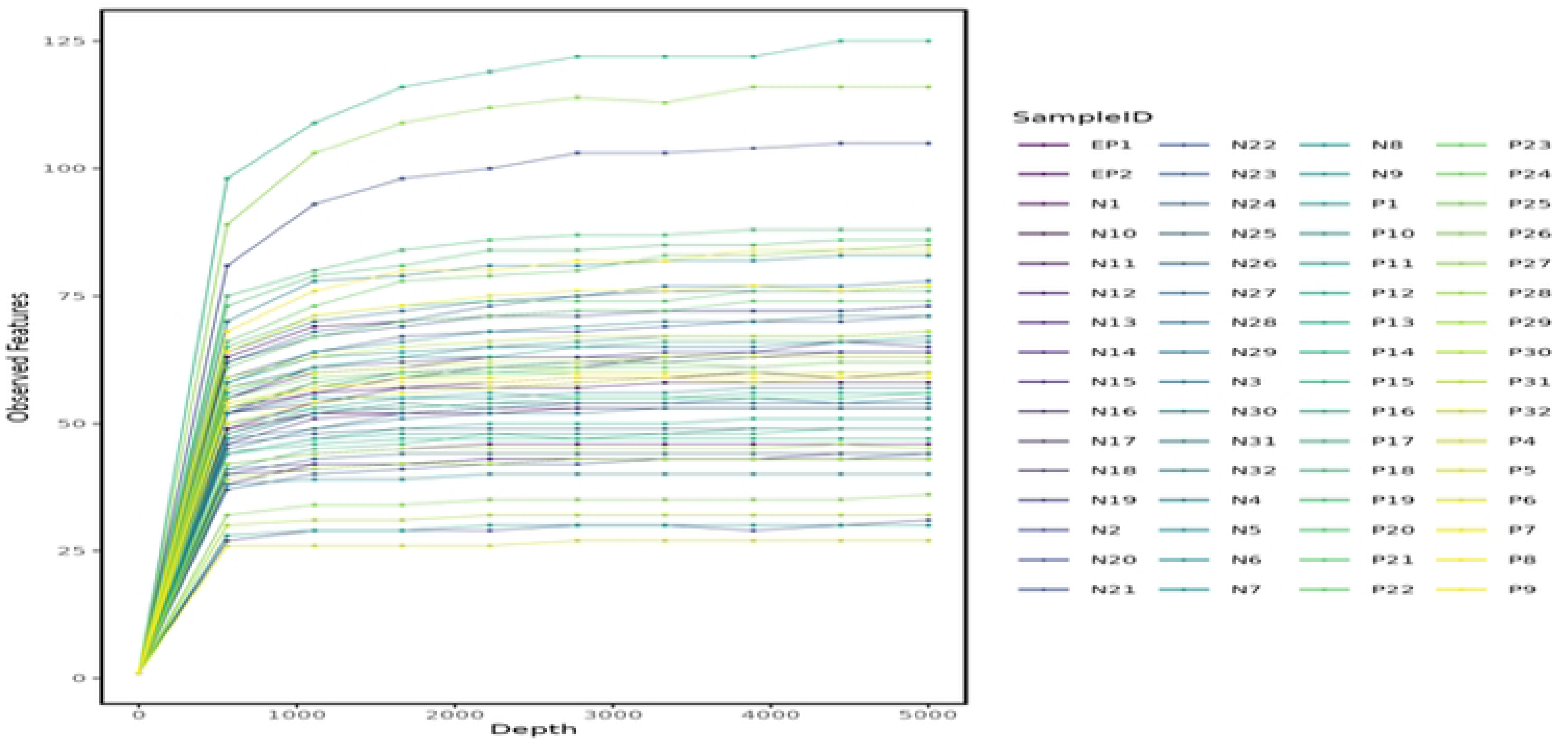
Rarefaction curve comparing COVID-19 positive (n=32) and negative (n=32) nasopharyngeal samples. Rarefaction data was obtained for all the study samples considering the depth of 0 to 5,000 sequences. Random colors were assigned to samples to facilitate comparison between both the groups of samples.

**Table 2:** Determination of alpha diversity in nasopharyngeal samples.

| **Diversity Index** | **COVID-19-Negative**<br>**Mean ± (SD)** | **COVID-19-Positive**<br>**Mean ± (SD)** | **p value** |
| --- | --- | --- | --- |
| Shannon (H) | 2.44 ± (0.075) | 2.18 ± (0.08) | 0.532 |
| Simpson (D) | 0.61 ± (0.11) | 0.65 ± (0.12) | 0.501 |
| Observed Richness | 68.80 ± (28.40) | 56.12 ± (19.05) | 0.192 |
| Observed Evenness | 0.52 ± (0.11) | 0.51 ± (0.12) | 0.642 |

The heatmap analysis and phylogenetic tree were generated for COVID-19-negative and COVID-19-positive samples separately and in comparison of both the groups. These nasopharyngeal swab samples showed the maximum frequency of phyla like Fusobacteriota, Actinobacteriota, Bacteroidota, Firmicutes, Proteobacteria, Campilobacterota, and Patescibacteria for COVID-negative samples. At the same time, Fusobacteriota, Actinobacteriota, Campilobacterota, Bacteroidota, Firmicutes, Proteobacteria, and Patescibacteria along with the bacterial variety at each site were predominant in COVID-positive samples. Further, a comparative heatmap analysis of negative and positive samples presented a significant frequency of Firmicutes, Proteobacteria, Bacteroidota, Actinobacteriota, Fusobacteriota, Patescibacteria and Campilobacterota as depicted in Figure 6A, 6B, 6C. The Relative frequency of occurrence of bacteria at taxonomic level was determined for COVID-19-positive (n=32) and -negative (n=32) nasopharyngeal samples. An average frequency of sequences per taxa was calculated from the total sum of all sequence counts, and the standard deviation (SD) is shown in percentage (%). The taxonomic data is included for phylum where its occurrence is > 1%, as depicted in Table 3. Furthermore, the compositions to determine the diversity of 16S rRNA metagenome for COVID-negative and positive samples are explored using the Krona chart, as shown in Figure 7A and 7B, respectively.

**Table 3:** Frequency of occurrence of bacteria at taxonomic level in COVID-19-positive and -negative nasopharyngeal samples.

| Phylum | Genus | COVID Negative<br>Avg.Frequency % ± (SD) % | COVID Positive<br>Avg.Frequency % ± (SD) % |
| --- | --- | --- | --- |
| Firmicutes | *Veillonella* | 22.36 ± (10.38) | 27.54 ± (12.46) |
| Firmicutes | *Streptococcus* | 6.67± (6.49) | 5.36± (3.46) |
| Firmicutes | *Megasphaera* | 1.32± (1.53) | 1.47± (1.86) |
| Proteobacteria | *Haemophilus* | 7.50± (6.30) | 6.33± (6.56) |
| Proteobacteria | *Neisseria* | 9.63± (8.74) | 10.39± (9.90) |
| Proteobacteria | *Actinobacillus* | 1.74± (4.73) | 1.23± (2.22) |
| Fusobacteriota | *Leptotrichia* | 1.50± (2.13) | 2.51± (3.26) |
| Fusobacteriota | *Fusobacterium* | 3.57± (2.97) | 3.52± (3.27) |
| Fusobacteriota | *Leptotrichia* | 1.34± (1.63) | 1.25± (2.35) |
| Bacteroidota | *Prevotella* | 2.48± (3.69) | 5.57± (4.47) |
| Bacteroidota | *Prevotella* | 6.12 ± (5.75) | 2.85 ± (4.92) |
| Bacteroidota | *Prevotella* | 1.12 ± (1.33) | 2.00 ± (2.89) |
| Bacteroidota | *Prevotella* | 1.98± (2.26) | 1.00± (1.29) |
| Bacteroidota | *Capnocytophaga* | 1.10± (2.40) | 1.54± (4.89) |
| Bacteroidota | *Porphyromonas* | 1.03± (1.81) | 0.95± (1.76) |
| Bacteroidota | *Porphyromonas* | 1.08± (2.03) | 0.85± (1.24) |
| Actinobacteriota | *Actinomyces* | 1.18± (1.63) | 1.17± (1.42) |
| Actinobacteriota | *Blastococcus* | 1.02 ± (2.74) | 0.68 ± (1.42) |
| Actinobacteriota | *Corynebacterium* | 1.99 ± (6.90) | 0.52± (0.93) |
| Patescibacteria | TM7x | 1.21 ± (1.53) | 1.23 ± (1.37) |

Overall, NGS analysis was done for all 64 study samples, considering 16S rRNA metagenomes of 32 COVID-positive and 32 COVID-negative nasopharyngeal swab samples. The outcome of this data, with the observation of variable frequency of phyla, genus, and species levels, was analyzed to understand the relative abundance of the bacteriome at the taxonomic level.

## Conclusion

In conclusion, the overall viral infections are considerably higher among SARS-CoV-2 negative patients (37.5% Vs 6.25%) as compared to SARS-CoV-2-positive patients representing respiratory illness probably due to interference of viral entities as well as competition benefit of SARS-CoV-2 in altering the imperviousness of the host. This pilot-scale study helped understand the significance of symptoms in SARS-CoV-2-negative individuals and the possibility of other viral infections. However, we were limited by the number of samples and follow-up of the patient to observe the later course of disease and severity. A well-funded extensive study with more samples from different locations and provision for long-term follow-up could shed more light on the significance of other infections with overlapping symptoms and co-infection impact in SARS-CoV2 positive patients.

Respiratory viral pathogens and bacteriome are unusually diverse, especially in SARS-CoV-2 infections. The metagenomic analysis explored that SARS-CoV-2 infection significantly impacted the diversity and the existence of a significant amount of bacteriome and respiratory viral pathogens in symptomatic COVID-positive individuals. The analyzed parameters of microbiome diversity identified in this study may be associated with the improvement, cure and resolution for COVID-19-positive individuals.

## Acknowledgments

All authors acknowledge ICMR-HQ and ICMR-NIV for their financial and material support as well as cooperation during the conduct of this work respectively. The authors also acknowledge the support of all the staff members who helped directly or indirectly with the collection and processing of these samples.

## Supporting information

### Authors’ contributions

VN: Conceptualization, Supervision, and Manuscript finalization, and communication

SJ: Review of Literature and Manuscript Writing, Sample processing for assay

RBW: Review of Literature and Manuscript Writing, assay performance

VAP: Facilitated the assays, analysis, and interpretation of data, manuscript review, and approval.

MLC: Facilitated the assays, analysis, and interpretation of data, manuscript review and approval

### Consent for publication

The study describes no individual data that may identify the individual. At the same time, the study had the waiver of consent by the institutional ethics committee and the approval for publication was obtained from the Research Integrity Unit of ICMR-NARI through the Director of the institute.

### Availability of data and materials

Most of the relevant data is being presented in the manuscript and other details are available with the corresponding author and would be provided on demand.

The sequencing database can be accessed using following details:

Submission ID: SUB14328959, BioProject ID: PRJNA1092386, https://www.ncbi.nlm.nih.gov/sra/PRJNA1092386

### Funding

The study was funded institutionally and no extramural funds were obtained for the same.

